# The Insulin receptor and Insulin like growth factor receptor 5’ UTRs support translation initiation independently of eIF4G1

**DOI:** 10.1101/2022.07.26.501659

**Authors:** Nicholas K. Clark, Meghan T. Harris, Michael T. Marr

**Author notes:** To whom correspondence should be addressed: Michael T. Marr II.

## Abstract

Protein synthesis is tightly regulated under stress conditions where energy may be scarce. Despite global repression of translation, some transcripts remain actively translated in order for the cell to be able to respond to the insult or prepare to quickly return normal cellular function after the stress ends. For the insulin receptor (Insr) and insulin-like growth factor receptor (Igf1r) transcripts this translation is mediated by an internal ribosome entry site (IRES) in their 5’UTRs that functions independently of eukaryotic initiation factor 4A (eIF4A) and eIF4E. Here we show that these cellular IRESes are also able to promote translation independently of the scaffolding protein eIF4G1 both *in vitro* and in the cell.

**Background:** IRES mediated translation initiation requires a different repertoire of factors than canonical cap-dependent translation.

**Results:** Treatments that inhibit the canonical translation factor eIF4G1 have little or no effect on the ability of the Insr and Igf1r cellular IRESes to promote translation.

**Conclusion:** Transcripts for two cellular receptors contain RNA elements that facilitate translation initiation without intact eIF4G1

**Significance:** Cellular IRES mechanisms may resemble viral type III IRESes allowing them to promote translate with a limited number of initiation factors allowing them to work under stress conditions when canonical translation is repressed.

## INTRODUCTION

Eukaryotic mRNA translation initiation requires a number of factors in order to facilitate recruitment of the ribosome to the 5’ end of the mRNA. The 5’ 7-methyl guanine (m^7^G) cap is bound by the cap binding complex, eukaryotic initiation factor (eIF) 4F, which is composed of three subunits: eIF4E, eIF4A, and eIF4G1[1]. eIF4E is the cap binding protein which directly binds to the m^7^G cap and recruits the other members of the eIF4F complex through its interaction with eIF4G1[1]. Because eIF4E is the lowest abundance subunit in the eIF4F complex, cap binding is thought to be the rate limiting step of canonical cap-dependent translation[2, 3]. eIF4A is a member of the DEAD-box family of helicases and uses ATP to unwind secondary structure in the 5’ untranslated region (5’ UTR) of the mRNA in order to facilitate ribosome binding[4, 5].

The canonical eIF4F complex is held together by the scaffolding protein eIF4G1 which contains binding sites for the other members of the eIF4F complex. eIF4G1 also facilitates recruitment of the small subunit of the ribosome through a direct interaction with eIF3[6-8]. The eIF3 complex contains 13 subunits and, together with the eIF2-Met-tRNA_i_^Met^-GTP ternary complex and the 40S ribosomal subunit make up the 43S pre-initiation complex[9-11]. Artificially recruiting the core domain of eIF4G1 to an mRNA is sufficient for recruitment of the ribosome and translation initiation highlighting the central role of this protein in initiation [12].

Internal ribosome entry sites (IRESes) were originally discovered in viruses as an alternative translation initiation mechanism[13, 14]. IRES elements allow for translation initiation independent of the m^7^G cap which is required for canonical cap-dependent translation. This allows transcripts containing IRES elements to be translated under conditions where canonical translation is repressed by inhibitors such as eIF4E binding protein (4E-BP) [15]. The IRES itself is a cis-acting RNA element where activity is generally mediated by RNA secondary structure[16, 17]. This RNA structure allows the UTR of the mRNA to bind directly to components of the translation initiation machinery independently of m^7^G cap binding by eIF4E.

Recently, viral IRESes have been categorized into eight classes depending on the minimal initiation factors required for translation initiation to occur[18]. IRESes in the *Dicistroviridae* family of viruses, exemplified by Cricket paralysis virus (CrPV), are the simplest. They require no translation initiation factors as they are able to directly bind to the A site of the 40S ribosomal to initiate translation. IRESes in the *Flaviviridae* family of viruses, the most well-studied of which is Hepatitis C virus (HCV) require eIF3 and a ternary complex, such as eIF2-Met-tRNA_i_^Met^-GTP in order to initiate translation. The largest class of identified viral IRESes are contained within the picornavirus family of viruses. These IRESes not only have the highest requirement for initiation factors but also make use of other IRES trans-acting factors (ITAFs).

While the factor requirements and mechanisms that viral IRES elements employ to engage the ribosome have been well studied, IRES elements in cellular mRNAs are much more poorly understood and do not seem to be as easily classified as their viral counterparts[18, 19]. This may in part be due to the situational nature of cellular IRES activity. Unlike some viral IRESes these cellular transcripts contain a ubiquitous m^7^G cap and consequently, they can be translated in a canonical cap-dependent manner. The IRES activity becomes important under conditions when the canonical pathway is inhibited such as cellular stress. Many identified cellular IRESes are found in transcripts that encode proteins important under these cellular conditions.

We previously identified cellular IRES elements in transcripts encoding proteins important for the insulin response in both *Drosophila* and mammalian cells [20-22]. The insulin receptor (Insr) and insulin-like growth factor receptor (Igf1r) transcripts of *M. musculus* contain IRES elements and are able to support cap-independent translation[20]. This was shown both *in vitro* using excess m^7^G cap analogue to inhibit the cap-binding protein, and in cells using bicistronic reporter constructs where second open reading frame (ORF) translation is dependent on IRES activity. We have also previously shown that these cellular IRESes are resistant to inhibition of the eIF4A helicase by the small molecule inhibitor hippuristanol [20, 23].

Here we show that translation driven by the Insr and Igf1r 5’UTRs is highly resistant to inhibition or depletion of the canonical translation factor eIF4G1. Taken together these results indicate a non-canonical mechanism for translation initiation of the Insr and Igf1r transcripts and support the idea that these transcripts utilize internal ribosome entry for translation initiation. The eIF4G1 scaffold is required for both canonical cap-dependent translation as well as translation driven by several classes of picornavirus IRESes. The findings described here suggest that the factor requirements for translation initiation driven by the Insr and Igf1r 5’UTRs may more closely resemble those of class III viral IRESes or may represent a distinct mechanism altogether [18, 19]. Further work on these important transcripts is necessary to resolve this question.

### EXPERIMENTAL PROCEDURES

#### Purification of his-tagged 2A-protease and eIF4G1^(1015-1118)^

Poliovirus 2A-protease was cloned into pMtac, an *E. coli* expression vector, with an N-terminal 6x His tag using PCR and standard cloning methods. eIF4G1^(1015-1118)^ was cloned into the pET28 *E. coli* expression vector with an N-and C-terminal 6x His tag using PCR and standard cloning methods. The plasmids were transformed into the BL21*(DE3) *E. coli* strain (Invitrogen) containing the pLacIRARE2 plasmid (Novagen). Cultures were grown to ∼0.5 OD600 and induced with 1mM IPTG overnight at 25°C. eIF4G1^(1015-1118)^ was purified using Ni Sepharose 6 Fast Flow (GE Healthcare) according to manufacturer’s instructions and eluted using 500mM imidazole. 2A-protease was purified using the same procedure with the addition of 0.1% sarkosyl during protein extraction to solubilize the 2A protein. Both proteins were dialyzed into storage buffer: 50mM Tris pH7.5 150mM NaCl 10% Glycerol; and snap frozen and stored at -80°C.

#### Immunobot of eIF4G1

For *in vitro* experiments 6 µL RRL treated with 1 µL purified 2A-protease were loaded onto a 10% SDS-PAGE gel. For cell-based assays 40µg total protein were used. Separated proteins were transferred to a nitrocellulose membrane. Blots were probed with mouse anti-eIF4G11 monoclonal antibody (clone2A9, Sigma) (1:1000) and imaged using Dylight secondary antibodies (Thermo). For cell-based assays, membranes were probed with the following antibodies: anti-EIF4G rabbit monoclonal antibody (1:5000, C45A4, Cell Signaling), mouse monoclonal Anti-HA (clone 12CA5), mouse E7 α-Tubulin antibody (1:500, DSHB), eIF4E Rabbit mAb (1:1000, C46H6, Cell Signaling) eIF4A rabbit mAb (1:1000, C32B4, Cell Signaling) β-Actin Mouse mAb (1:1000, 8H10D10, Cell Signaling) and detected using Dylight infrared secondary antibodies (Thermo) on a ChemiDocMP imager (Biorad).

#### In vitro transcription and RNA reporter preparation

Transcription templates for RNA reporters were prepared by digesting plasmid templates containing the UTRs of interest upstream of a firefly luciferase ORF all under the control of a T7 or T3 reporter. Templates were purified using a column DNA clean up protocol (Biobasic) and eluted in TE pH 8.0. Templates were transcribed using T3 or T7 RNA polymerase for 4 hours at 37°C and purified using LiCl precipitation. A 5’ 7meG cap was added to the RNA reporters using vaccinia virus capping enzyme (NEB) according to the manufacturer’s protocol. Capped reporters were purified using phenol/chloroform extraction and isopropanol precipitation. RNAs were then 3’ poly(A) tailed using *E. coli* poly(A) polymerase (NEB) and purified using phenol/chloroform extraction and isopropanol precipitation.

#### In vitro Translation

Translation assays were done in a 10 µL reaction volume using 6 µL of whole rabbit reticulocyte lysate still containing endogenous mRNAs to allow for competitive conditions. Translation extracts were pre-incubated with 130 nM eIF4G1^(1015-1118)^ for 10 minutes at 30°C, 2A-protease for 1.5 hours at 30°C, or the respective buffer control. Translation reactions also contained: 24 mM HEPES pH 7.4, 0.4 mM magnesium acetate, 30mM potassium acetate, 16.8 mM creatine phosphate, 0.1mM spermidine, 60 µM Amino Acids, 800 ng creatine kinase, 1 µg calf liver tRNA, and 100 ng of reporter RNA. Translations reactions were incubated at 37°C for 30 minutes. Luciferase activity was measured using 100 µL of luciferase substrate: 75 mM HEPES pH8.0, 5 mM MgSO_4_, 20 mM DTT, 530 µM ATP, 100 µM EDTA, 500 µM coenzyme A, and 500 µM D-luciferin.

#### Cell Culture, transient transfections, bicistronic dual luciferase assay

HEK293T and HCT116 cells were maintained in Dulbecco’s Modified Eagle’s Medium supplemented with 10% FetalPlex (Gemini Bio), 1x Penicillin/Streptomycin, 1 µg/mL Insulin. For transient transfection cells were seeded in a 24-well plate at a density to be ∼60% confluent at the time of transfection and all transfections were done with a 1:5 bicstronic reporter to expression ratio. HEK293T cells were transfected with bicistronic reporter plasmids and either CMV driven eIF4G1^(1015-1118)^, or an empty FLAG-tag expression vectors with polyethylenimine at a 3:1 PEI to DNA ratio. eIF4G1^(1015-1118)^ experiments were harvested 48 hours after transfection with Passive Lysis Buffer (Promega). HCT116 cells were transfected with jetPRIME (Polyplus) according to manufactures protocol. CMV driven eIF4G1^(1015-1118)^ transfections were harvested after 72 hours. Firefly and renilla activity were measured using the dual-luciferase reporter system (Promega). Firefly luciferase was measured in 75 mM HEPES pH8.0, 5 mM MgSO_4_, 20 mM DTT, 530 µM ATP, 100 µM EDTA, 500 µM coenzyme A, and 500 µM D-luciferin. Renilla luficerase was measured by adding an equal volume of 1.0 M NaCl, 500 mM Na_2_SO_4_, 25 mM Na_4_PPi, 15 mM EDTA, 10 mM NaOAc, and 0.1 mM coelenterazine.

#### Mammalian Cell Culture

HCT116 (human colon carcinoma) cells were maintained in Dulbecco’s Modified Eagle Medium (Genesee Scientific) supplemented with 10% FetalPlex (Gemini Bio), Penicillin-Streptomycin (Genesee Scientific), and Bovine Insulin (MilliporeSigma). HCT116 cells were seeded at a density of 8 × 10^4^ cells per milliliter for a 72 hour total incubation period, transfected with JetPrime (Polyplus), and maintained with a subculture ratio of 1:6.

#### Generation of eIF4G1 dTAG line

eIF4G1 Targeting constructs were based on plasmid pCRIS-PITChv2-dTAG-BSD (BRD4) a gift from James Bradner & Behnam Nabet (Addgene plasmid # 91795) [24] This construct contains the FKBP_F36V-2xHA-P2A-BSD (dTAG) cassette encoding, in frame, the FKBP_F36V protein followed by two copies of the HA epitope, a 2A peptide from porcine teschovirus-1 polyprotein and a blasticidin resistance marker. DNA corresponding to 271bp upstream of the eIF4G1I stop codon was amplified by PCR from HCT116 genomic DNA and inserted in frame with the dTAG coding sequence. In the PCR process, silent mutations were introduced in the coding region for the last five amino acids of eIF4G to prevent CRISPR/cas9 from targeting the recombinant locus. Subsequently, DNA corresponding to 449bp downstream of the stop codon was amplified by PCR from HCT116 genomic DNA and inserted after the dTAG cassette. A second construct containing the neomycin resistance marker in place of the blasticidin resistance marker was created.

A guide RNA expressing plasmid (phU6_eIF4G1_gRNA_2611) was created by inserting annealed, phosphorylated oligonucleotides (cacc<u>GAGGAGTCTGACCACAACTG</u>, aaac<u>CAGTTGTGGTCAGACTCCTC</u>) targeting the last 20 bases of eIF4G1 into the BbsI site of phU6, a derivative of pX330-U6-Chimeric_BB-CBh-hSpCas9 (a gift from Feng Zhang Addgene plasmid # 42230) lacking the cas9 expression cassette.

The neomycin and blasticidin targeting constructs were co-transfected into HCT116 cells using JetPrime (Polyplus) along with phU6_eIF4G1_gRNA_2611 and pJDS246 expressing cas9. pJDS246 was a gift from Keith Joung (Addgene plasmid # 43861). After 3 days the cells were treated with 30µg/ml blasticidin followed by 100µg/ml G418. Cell lines were cloned by picking resistant colonies followed by cloning by dilution. Cell lines were screened for insertion by PCR and by western. Positive lines were tested for eIF4G1 degradation using 1µM dTAG-13 and 1µM dTAGV-1.

#### eIF4G1 Degradation

To achieve eIF4G1 degradation in the HC116 C8 clonal line, 1uM dTagV-1 (Tocris Bioscience) was added in complete media and incubated for 1.5 hours at 37°C before any assays were conducted, including reporter transient transfections, puromycin incorporation, and polysome gradient centrifugation.

#### Polysome Gradient Centrifugation

11mL,10-50% sucrose gradients were prepared in Beckman polyallomer tubes (No. 331372). Gradients were assembled by layering 11 levels of 1mL sucrose solution with gradually decreasing concentrations, from 1.5M to 0.5M. These sucrose solutions were buffered using 50mM Tris PH7.5, 0.25M KCl, and 50mM MgCl_2_ and supplemented with 0.1mg/mL cycloheximide. Each sucrose layer was added with an 1mL serological pipet (with gravity mode on pipet aids) without disrupting the separating surface. Gradients were sealed with parafilm, stored at -80°C and thawed at 4°C overnight before use. Cells were incubated with 0.1mg/mL cycloheximide for 5 minutes at 37°C for 5 minutes. Cells were harvested in media, centrifuged at 800X g for 5 minutes, and washed with 1mL ice-cold PBS with 0.1mg/mL cycloheximide. Cells were centrifuged at 15Xg for 5 minutes at 4°C. Cell pellets were flash frozen in liquid nitrogen and stored overnight at -80°C. Cell pellets were lysed in 20mM HEPES pH7.4, 150KCl, 2.5mM MgOAc_2_, 1% Triton X-100, 1X EDTA-free protease inhibitors (Sigma), 1mM Dithiothreitol (DTT), 0.1mg/mL cycloheximide, and 0.1 μL/mL superasin (Life Technologies) for 10 minutes on ice and centrifuged at 21000g for 10 minutes at 4°C. 15 A260 units of induced and uninduced extracts were loaded onto 11mL sucrose gradients described previously. Sucrose gradients were centrifuged for 2 hours at 35000 rpm in a SW41 Ti rotor in a Beckman ultracentrifuge. Gradients were then collected using a Brandel syringe pump and A254 was measured using a Teledyne Isco UA-6 absorbance detector.

#### Puromycin Incorporation Assay

Cells were treated with 1µM dTagV-1 as indicated. For cycloheximide treated samples, cells were treated with 0.1mg/mL cycloheximide for 5 minutes at 37°C. All samples were then incubated with 10µg/mL puromycin for 30 minutes at 37°C. Cells were harvested in 1X RIPA buffer (20 mM Tris-HCl (pH 7.5), 150 mM NaCl, 1 mM Na_2_EDTA, 1 mM EGTA, 1% NP-40, 1% sodium deoxycholate, 2.5 mM sodium pyrophosphate, 1 mM β-glycerophosphate, 1 mM Na_3_VO_4_, and 1µg/mL leupeptin with 1X EDTA-free protease inhibitors (Sigma)). Protein concentrations were determined using BCA assay (Pierce) and equal amounts of protein were loaded onto a 4-12% SDS-PAGE gel. Protein was transferred to an activated (PVDF) membrane, blocked, and probed with either mouse anti-puromycin antibody PMY:2A4 (1:1000, DSHB) and mouse E7 α-Tubulin antibody (1:500, DSHB). The anti-puromycin PMY:2A4 antibody has been described[25]. The E7 tubulin antibody was developed by M. Klymkowsky at the University of Colorado/MCD Biology. Both antibodies were obtained from the Developmental Studies Hybridoma Bank, created by the NICHD of the NIH and maintained at The University of Iowa, Department of Biology, Iowa City, IA 52242.

## RESULTS

### Insr and Igf1r cellular IRES-mediated translation is stimulated by eIF4G1 cleavage with poliovirus 2A-protease

To begin to examine the eIF4G1 requirement of these two cellular IRESes we started by using 2A-protease, a natural eIF4G1 targeting system from poliovirus. Recombinant poliovirus 2A-protease specifically cleaves eIF4G1, separating the N-terminal domains containing the eIF4E and PABP binding sites from the C-terminal domain, which binds to RNA, eIF4A and eIF3 (Fig. 1A)[26]. We expressed the poliovirus 2A-protease as a 6-histidine fusion protein in *E. coli*. The 2A protease fusion protein was purified using metal affinity chromatography (Fig. 1B).

**Figure 1.**
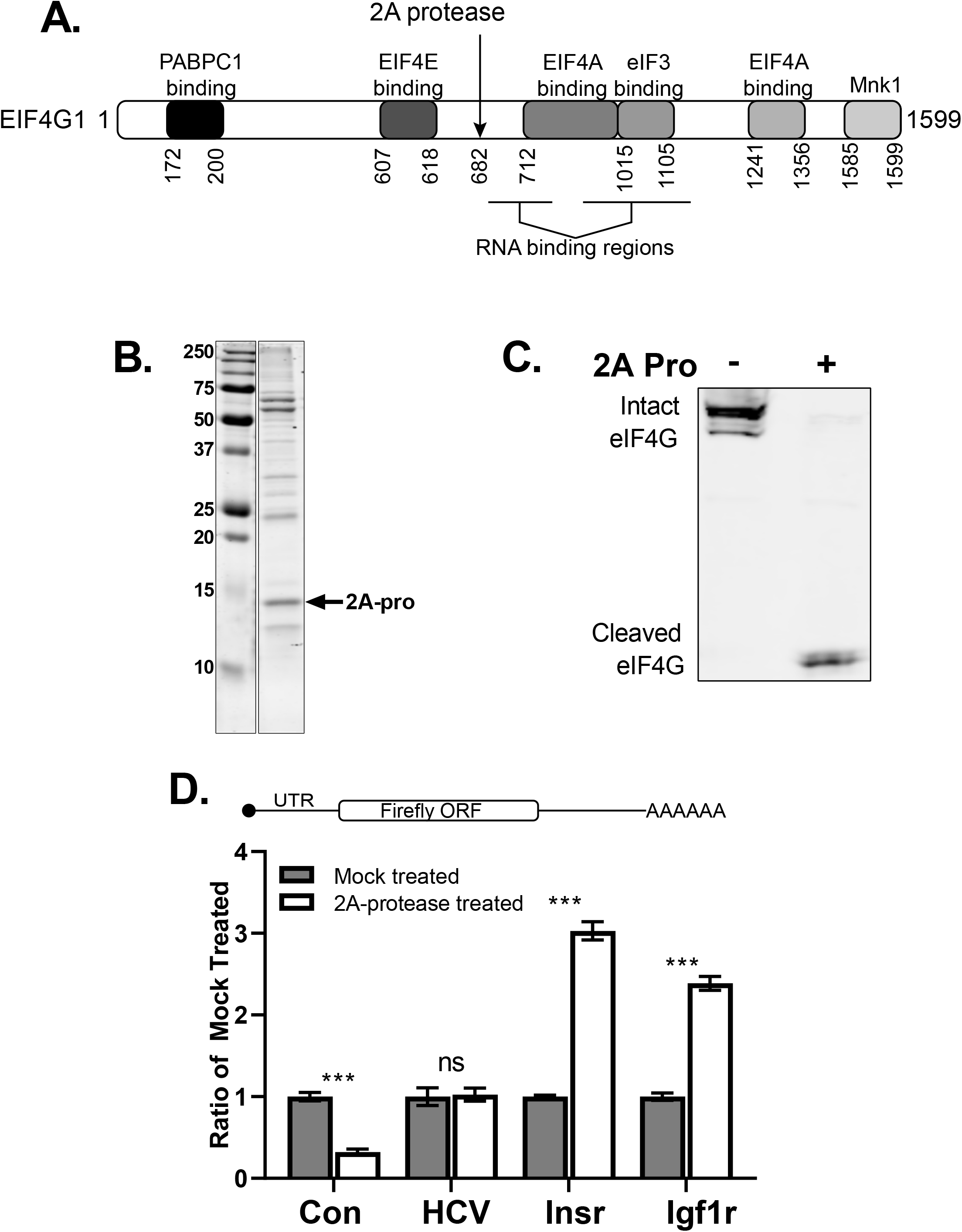
Insr and Igf1r cellular IRESes are resistant to eIF4G1 cleavage by poliovirus 2A-protease *in vitro*. **A**. Schematic of binding domains on eIF4G1. The 2A-protease cleavage sight is marked with an arrow. **B**. SDS-PAGE of recombinant 2A-protease purified from *E. coli*. **C**. Western blot showing purified 2A-protease is able to cleave eIF4G1 in RRL translation extracts. **D. (top)** Diagram of RNA reporters used in the *in vitro* translation assays. **(bottom)** Activity of these RNA reporters in RRL *in vitro* translation extracts (37°C 30 min) treated with poliovirus 2A-protease (30°C 1.5 hours) Data are plotted as the fraction of the respective buffer treated control. (error bars indicate SEM, * represents p-value ≤ 0.05, ** represents p-value ≤ 0.01, *** represents p-value ≤0.001)

Purified 2A protease was used to treat Rabbit Reticulocyte Lysate (RRL) translation extracts to cleave the eIF4G1 present in the extract. We assayed the extent of eIF4G1 cleavage by immunoblotting with a monoclonal antibody raised against amino acids 546-645 of eIF4G1 capable of detecting the amino-terminal fragment of the cleaved eIF4G1. Using this assay, we determined the optimum amount of 2A required for maximum eIF4G1 cleavage. With the optimized conditions, almost all the detectable eIF4G1 is cleaved (Fig. 1C).

We then programmed the 2A-protease-treated RRL or mock-treated RRL translation extracts with *in vitro* transcribed, capped and polyadenylated firefly luciferase monocistronic mRNA reporters. mRNAs containing the 5’ untranslated region (UTR) of mouse Insr, mouse Igf1r, Hepatitis C Virus, or a cap-dependent reporter were assayed (Fig. 1D). Translation activity was measured by assaying the firefly luciferase protein produced in 30 minutes at 37°C. In the 2A-protease-treated extracts cap-dependent translation is strongly inhibited (Fig. 1D). The HCV viral IRES reporter is unaffected by 2A-protease-treatment, indicating that the extract is not generally inhibited and that the deficiency is specifically in cap-dependent translation. Translation of the RNAs containing the Insr and Igf1r UTRs is not inhibited by eIF4G1 cleavage. In fact, the translation activity of RNAs containing the Insr and Igf1r UTRs is stimulated compared to mock-treated extracts (Fig. 1D). This result is consistent with previous experiments that showed Insr and Igf1r IRES containing reporters, as well as the viral IRES containing from HCV or EMCV, have a competitive advantage in non-micrococcal nuclease treated extracts when cap-dependent translation is inhibited[20, 27].

### Insr *and* Igf1r *cellular IRES mediated translation is resistant to competitive inhibition by the eIF4G1-eIF3 binding domain*

2A protease cleavage of eIF4G1 separates the eIF4E binding region from the eIF3 binding region. However, as seen with the poliovirus IRES, the carboxy terminus of eIF4G1 can function to recruit eIF3 and the 40S subunit by interacting directly with the IRES. To investigate this mechanism for these cellular IRESes, we purified a fragment of eIF4G1 (amino acids 1015-1118) which contains the eIF3 binding site and used it as a competitive inhibitor (Fig. 2A). Previous work has shown that the eIF4G1/eIF3 interaction is mediated by a ∼93 amino acid region of eIF4G1[6, 7, 28]. Importantly, this binding domain alone can compete with full length eIF4G1 for eIF3 binding and functionally disrupt this interaction[8].

**Figure 2.**
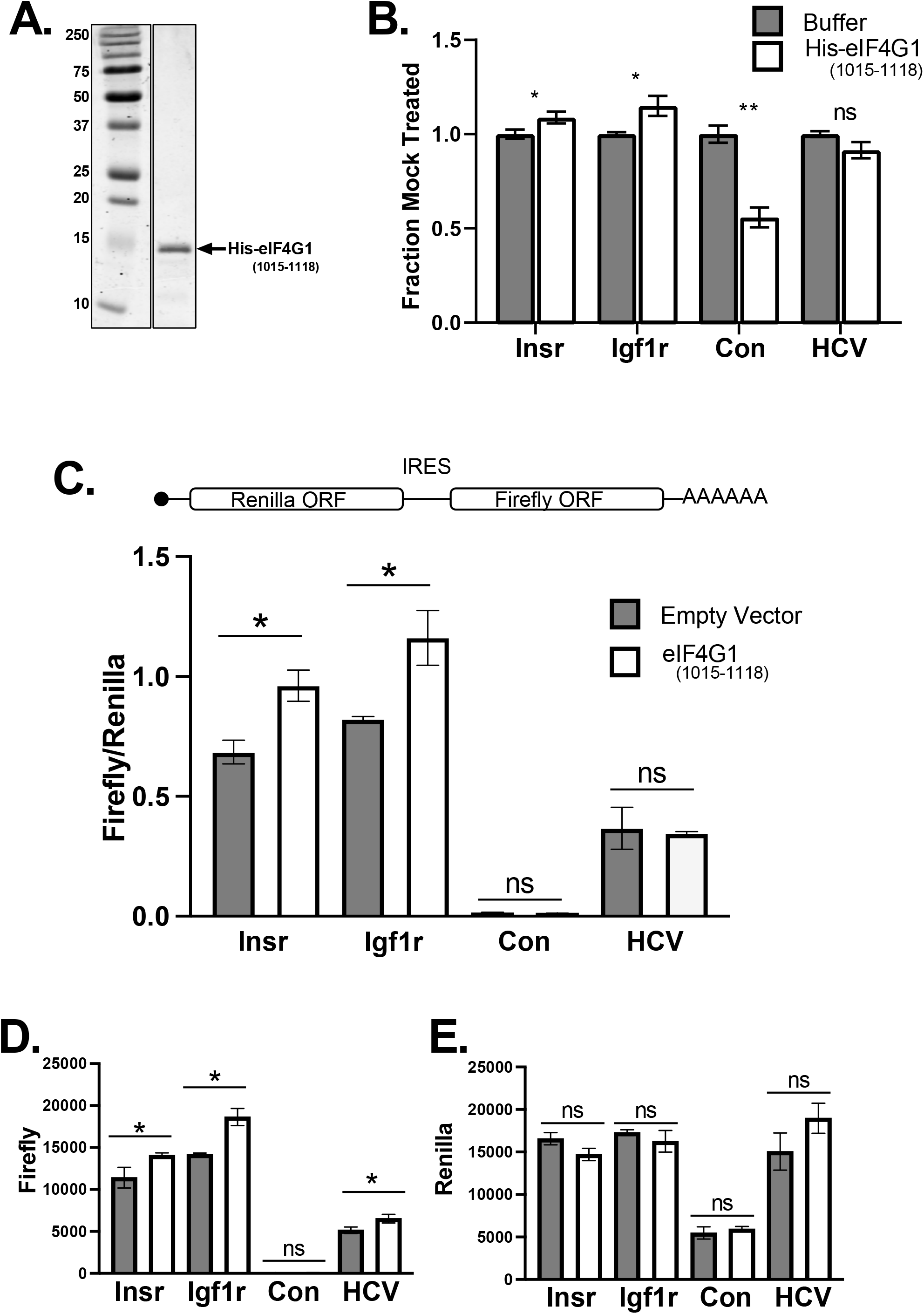
Insr and Igf1r cellular IRESes are resistant to competitive inhibition of eIF4G1/eIF3 interaction by eIF4G1(1015-1118) fragment. **A**. SDS-PAGE of recombinant eIF4G1(1015-1118) fragment purified from *E. coli*. **B**. Activity of these RNA reporters in RRL *in vitro* translation extracts (37°C 30 min) with and without 130 nM eIF4G1(1015-1118) fragment. Data are plotted as the fraction of the respective buffer treated control. **C**. IRES activity of transiently transfected bicistronic reporters shown as firefly to renilla ratio. Reporters were cotransfected with an empty vector control or a vector expressing the eIF4G1(1015-1118) fragment. **D**. Firefly luciferase activity of these reporters. **E**. Renilla luciferase activity of these reporters. (error bars indicate SD, * represents p-value ≤ 0.05, ** represents p-value ≤ 0.01,)

RRL translation extracts were supplemented with an excess of purified eIF4G1^(1015-1118)^ and again programed with the firefly luciferase monocistronic mRNA reporters. The cap-dependent mRNA reporter is inhibited approximately 50% by excess eIF4G1^(1015-1118)^. We can conclude that the fragment is not generally detrimental to the extract because the HCV viral IRES, which does not require eIF4G1, is unaffected by the excess eIF4G1 fragment (Fig. 2C). In these same conditions the Insr and Igf1r cellular IRESes were uninhibited. In fact, there is a significant, although small, stimulation of the translation activity for the cellular IRES containing transcripts.

### The Insr and Igf1r cellular IRESes are resistant to inhibition by the eIF4G1^(1015-1118)^ fragment in mammalian cells

In order to test the eIF4G1 requirements for the Insr and Igf1r IRESes in a cellular context we employed our well characterized bicistronic reporter constructs that we have previously used to test eIF4E and eIF4A requirements[20]. In these constructs the UTR of interest is cloned between a renilla luciferase ORF and a firefly luciferase ORF. RNA expression is driven by the Rous sarcoma virus LTR promoter creating a single bicistronic mRNA that contains both open reading frames (Fig. 2C). Translation of renilla luciferase is cap-dependent while expression of firefly luciferase depends on IRES activity. These reporters were co-transfected with a construct that uses the CMV promoter to drive expression of eIF4G1^(1015-1118)^ or an empty vector control. Compared to the empty vector, co-expression of eIF4G1^(1015-1118)^ causes a stimulation of Insr and Igf1r IRES activity, measured by the increase in the ratio of firefly luciferase activity to renilla activity. (Fig. 2C). Under these conditions, expression of eIF4G1^(1015-1118)^ has little effect on renilla activity itself but the stimulation of IRES activity is evident in the increase of firefly reporter activity alone (Fig. 2D&E).

### The Insr and Igf1r cellular IRESes promote translation initiation in the absence of eIF4G1 mammalian cells

To determine the absolute requirement for eIF4G1 in translation initiation mediated by the Insr and Igf1r IRESes we sought to rapidly remove eIF4G1 from the cell. We chose to use targeted degradation based on the dTAG system [24, 29]. This system utilizes FKBP12^F36V^ fusion proteins and a heterobifunctional small molecule to specifically degrade target proteins. A synthetic FKBP12^F36V^ ligand (AP1867) is connected to a small molecule that interacts with an E3 ligase, selectively targeting the fusion protein for degradation. We designed a fusion construct based on the dTAG system that would create an in-frame fusion with the C terminus of eIF4G1 (Fig. 3A). Using CRISPR/cas9 we recombined the construct into the genome of HCT116 cells and tested clones for cassette insertion and eIF4G1 degradation in response to the small molecule ligand.

**Figure 3.**
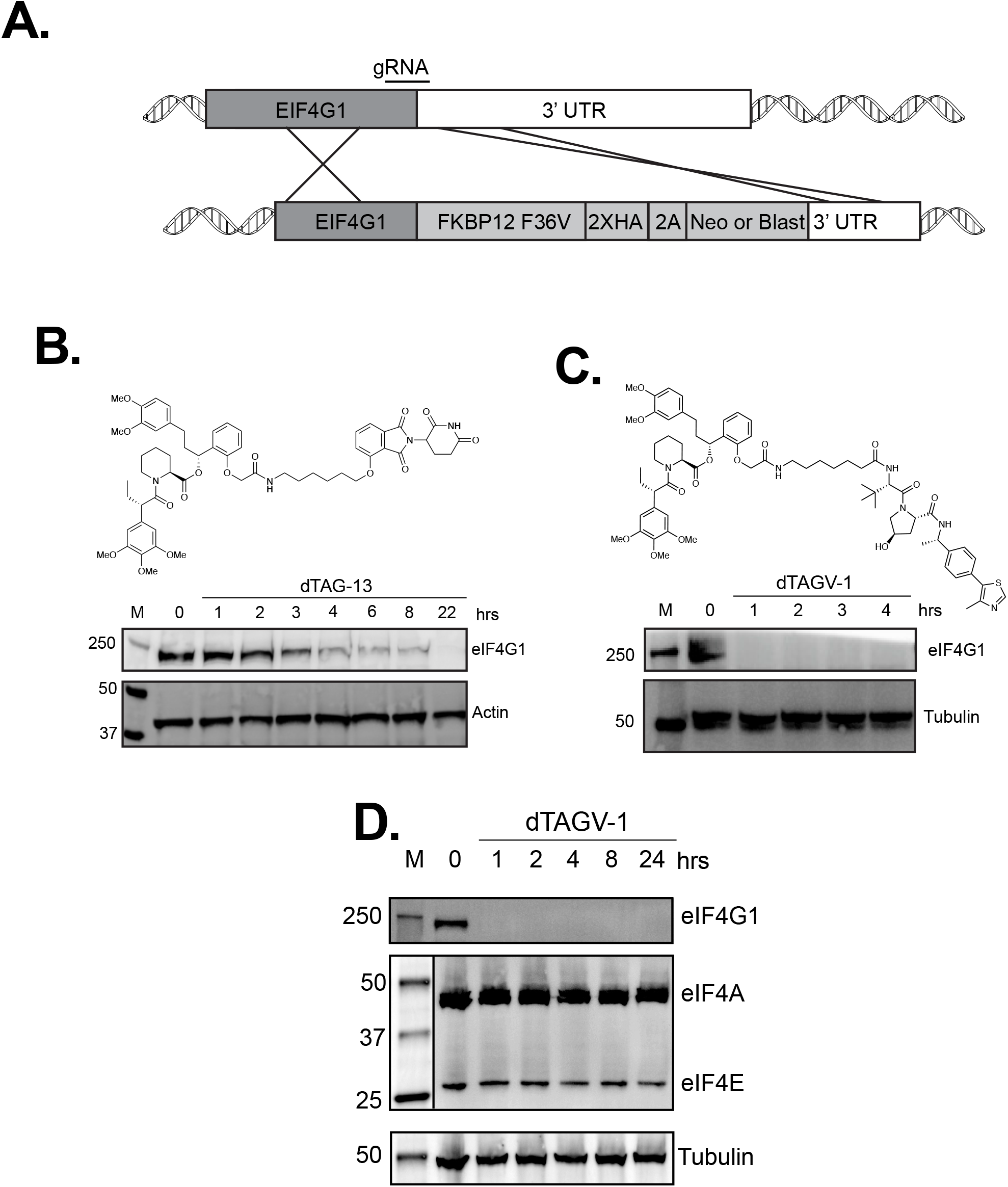
Creation of an eIF4G1 depletion cell line. **A**. Diagram of the targeting strategy for eIF4G1. **B**. Time course of treatment with 1µM dTAG-13 small molecule. eIF4G1 is detected by immunoblot, actin detection serves as a loading control. **C**. Time course of treatment with 1µM dTAGV-1 small molecule. eIF4G1 is detected by immunoblot, tubulin detection serves as a loading control. **D**. Time course of treatment with 1µM dTAGV-1 small molecule. eIF4G1, eIF4E and eIF4A are detected by immunoblot, tubulin detection serves as a loading control. In the middle panel the molecular weight marker is separated by a line to indicate it was detected in a different fluorescent channel.

We first tested the dTAG-13 small molecule. This molecule contains AP1867 attached to a ligand that binds the cereblon E3 ligase[24]. Cells were treated with 1µM dTAG-13 and samples were collected over a 24-hour time course. The loss of eIF4G1 was assayed by immunoblot. Addition of dTAG-13 was found to cause a significant loss of eIF4G1 from the cell with roughly 50% of the eIF4G1 is lost from the cell in 4 hours. However, more than 22 hours post treatment were required to see complete loss of eIF4G1 (Fig. 3B).

Next, we tested the dTAGV-1 small molecule. This molecule is AP1867 attached to a ligand that binds the von Hippel-Lindau (VHL) E3 ligase complex[30]. Cells were treated with 1µM dTAGV-1 and time course samples were collected to assay the loss of eIF4G1. Surprisingly, we observed that eIF4G1 is depleted in 1 hour (Fig. 3C). Quantitation of the eIF4G1 signal indicates a greater than 99% depletion at that timepoint. The combination of this cell line and dTAGV-1 provides a way to rapidly and specifically remove eIF4G1 from the cell. At the same time, we also monitored the other subunits of eIF4F to determine the dependence on eIF4G1 for their stability. There is no appreciable loss of either eIF4A or eIF4E during 24 hours of dTAGV-1 treatment in these cells (Fig. 3D).

The ability to rapidly deplete eIF4G1 allowed us to globally test the dependence of translation on eIF4G1 shortly after its degradation in this cell line. As 1 hour is sufficient to degrade the vast majority of eIF4G1, this rapid, specific degradation minimizes the possibility of confounding cellular compensation. We first looked at global translation using the puromycin incorporation assay. Cells were treated with dTAGV-1 for either 1.5 or 3 hours to deplete eIF4G1 (Fig. 4A). This should allow enough time for previously engaged ribosomes to run off their mRNA templates [31]. Cells were then pulse labeled with puromycin to detect proteins that are being actively translated. Total cellular protein was separated by SDS PAGE and puromycin incorporation was detected using a monoclonal antibody against puromycin[25]. Mock treated cells showed a robust incorporation of puromycin that depended on active translation as evidenced by its complete inhibition by cycloheximide (Fig. 4B). Upon eIF4G1 depletion puromycin incorporation is greatly decreased at 1.5 and 3 hours (67% and 80% respectively) indicating a global inhibition of translation (Fig 4B). However, there clearly remains detectable translational activity in eIF4G1 depleted cells.

**Figure 4.**
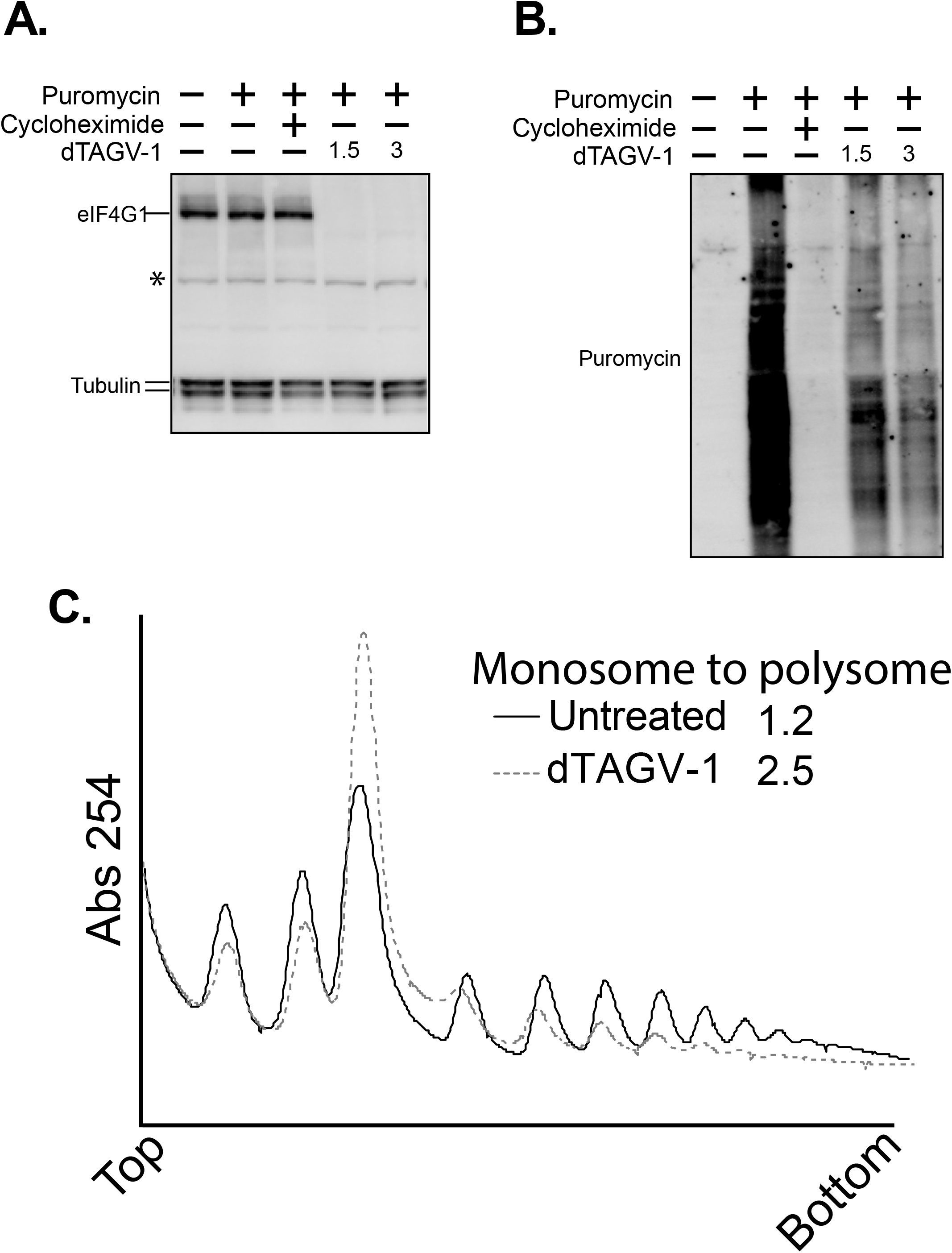
Global effects of eIF4G1 depletion. **A**. eIF4G1 is detected by immunoblot under the conditions tested for puromycin incorporation, tubulin detection serves as a loading control. **B**. The same membrane as in A but blotted with anti-puromycin antibodies and detected in a separate channel. **C**. Polysome profile of eIF4G1 dTAG line treated with dTAGV-1 for 1.5 hours.

To investigate the effect of eIF4G1 depletion on polysomes, cytoplasmic extracts were separated on a sucrose gradient and ribosomes were monitored by absorbance at 254nm. Comparison of mock treated cells with cells that have been treated with dTAGV-1 indicates a preferential loss of heavy polysomes. There is a concomitant increase in the population of monosomes in eIF4G1 depleted cells leading to a drastic increase in the monosome to polysome ratio (Fig. 4C).

We tested the activity of the Insr, Igf1r and HCV IRES in the eIF4G1 dTAG cell line using the bicistronic system described above. Cells were treated with dTAGV-1 for 1.5 hours to deplete endogenous eIF4G1. Under these conditions the cap-dependent renilla signal is reduced 50-60% (Fig. 5A, REN) for all reporters. The HCV IRES dependent firefly signal is similarly affected under these conditions, where it is reduced to 50% of that of untreated cells. By contrast, both the Insr and Igf1r IRES dependent firefly signals retain 80% activity in these conditions (Fig. 5A, FF).

**Figure 5.**
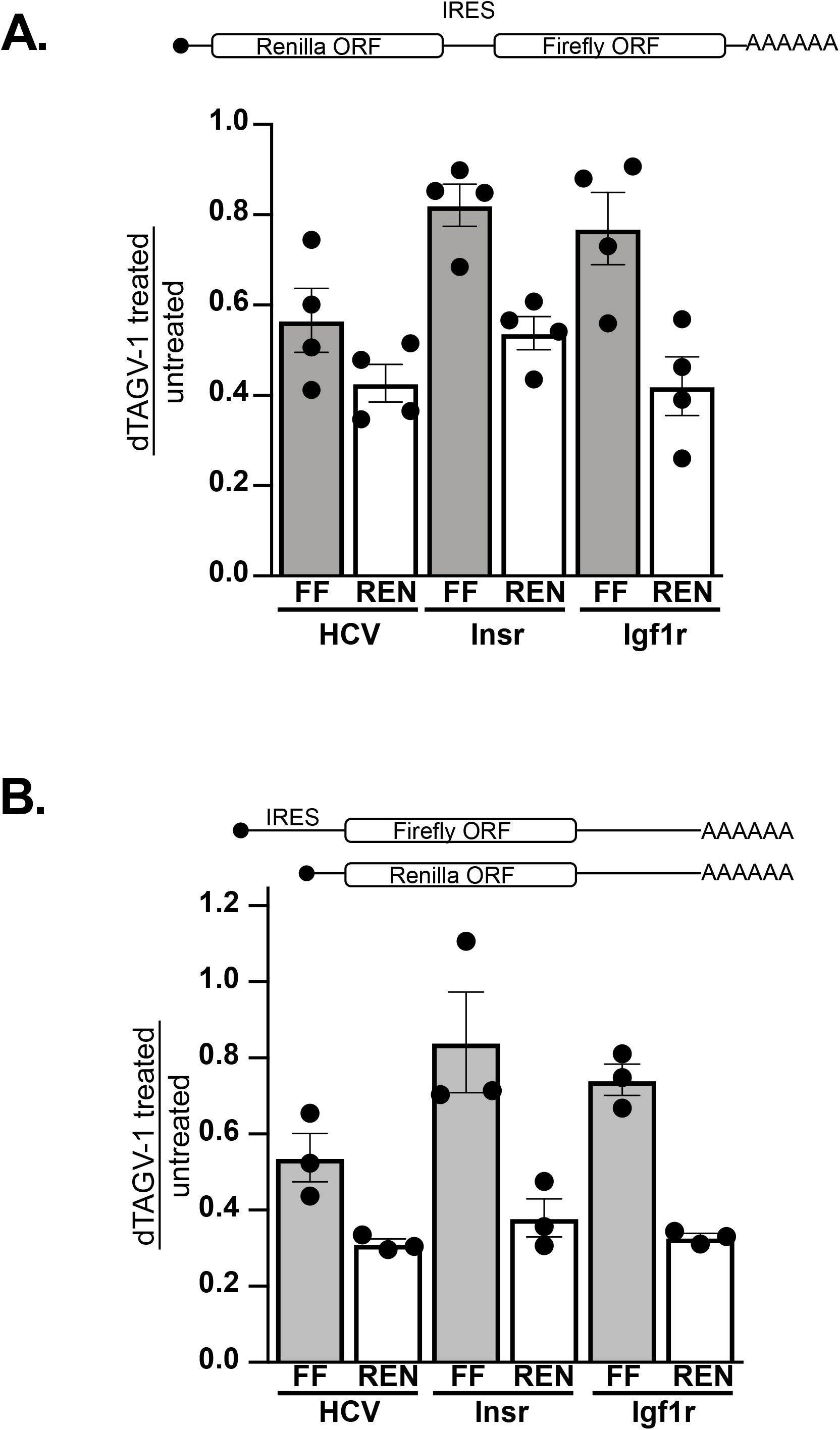
Effects of eIF4G1 depletion on cellular IRESes. **A. (top)** Diagram of bicistronic RNA produced by reporter constructs. **(bottom)** Activity of these reporters. Values are plotted as the ratio of dTAGV-1 treated cells to mock treated cells. FF indicates firefly luciferase activity. REN indicates renilla luciferase activity **B. (top)** Diagram of monocistronic RNAs produced by reporter constructs. **(bottom)** Activity of these reporters. Values are plotted as the ratio of dTAGV-1 treated samples to untreated. FF indicates firefly luciferase activity. REN indicated renilla luciferase activity. (error bars indicate SD, biological replicates are shown as filled circles)

To ensure this result is not a product of the bicistronic reporter system we repeated these experiments with monocistronic constructs (Fig. 5B). All IRES constructs were cotransfected with the same cap-dependent monocistronic renilla reporter. We found that the effect of eIF4G1 depletion on the monocistronic renilla construct to be more inhibitory as the cap-dependent translation reporter dropped to 40% when eIF4G1 was depleted in all transfections (Fig. 5B, REN). Again, the HCV IRES retains 50% activity in these depletion conditions and consistent with the bicistronic assay, the Insr and Igf1r cellular IRESes retain approximately 80% activity indicating a robust resilience to the depletion of eIF4G1 (Fig. 5B, FF).

## DISCUSSION

While the mechanisms of many viral IRESes are very well characterized[18], cellular IRES elements are much more poorly understood. Our lab has previously shown that the Insr and Igf1r transcripts contain IRES elements in their 5’ UTRs that are able to support translation initiation cap-independently *in vitro* and are able facilitate internal initiation on bicistronic reporter constructs *in vitro* and in cells. It has also been shown that these IRES are able to function independently of the initiation factor eIF4A[20]. However, the precise mechanisms by which these cellular IRES elements function are still not understood.

The translation factor eIF4G1 is critical for canonical cap-dependent translation and is also utilized by a number of viral IRESes for initiation[13, 32]. In addition, recruitment of eIF4G1 to an mRNA is sufficient to recruit the ribosome and initiate translation through its interaction with eIF3[12]. In order to test the requirement for eIF4G1 in translation initiation driven by the Insr and Igf1r cellular IRESes *in vitro* we perturbed eIF4G1 in two ways.

First, we utilized recombinant poliovirus 2A-protease which specifically cleaves eIF4G1 separating N-terminal eIF4E and PABP binding domains from the C-terminal eIF4A and eIF3 binding domains. Using RRL *in vitro* translation assays we show that cleavage of eIF4G1 actually stimulates translation of Insr and Igf1r reporter constructs, a result similar to one previously reported using *in vitro* cap competition experiments[20]. We believe that this stimulation represents an increased availability of translational machinery to the Insr and Igf1r reporters, as translation machinery has been re-tasked from mRNAs endogenous to the RRL extracts whose translation initiation is eIF4G1 dependent [27]. This result is consistent with the idea that the mRNAs containing the Insr and Igf1r UTRs support translation initiation under conditions of a compromised eIF4G1.

Second, we used a recombinant eIF4G1 fragment to competitively inhibit the eIF4G\eIF3 interaction that is thought to lead to ribosome recruitment. When eIF4G1^(1015-1118)^ is added to *in vitro* translation extracts cap-dependent translation is specifically inhibited while the Insr and Igf1r IRES remain uninhibited. When this same fragment is expressed in mammalian cells along with bicistronic IRES reporter constructs, the activity of both the Insr and Igf1r UTRs is stimulated as measured by the firefly/renilla ratio.

To investigate the eIF4G1 requirement of the Insr and Igf1r UTRs in cells, we created a cell line derived from the diploid HCT116 colon cancer line that allows the conditional depletion of eIFG1 using the dTAG system[24]. Utilizing the small molecule dTAG-V1 we are able to deplete eIF4G1 from the cell in 1 hour. Despite the depletion of the eIF4F scaffold, the levels of the other eIF4F components (eIF4E and eIF4A) are unchanged. eIF4G1 depletion results in an 80% decrease in active translation three hours after addition of the small molecule as measured by puromycin incorporation. Polysome profiles of the depleted cells show a preferential loss of heavy polysomes and an increase in monosomes consistent with an inhibition of translation initiation. The effects of eIF4G1 depletion on global translation, in this cellular context, are much more dramatic than previously reported [33]. Interestingly, the 20% residual translation correlates well with the predicted level of DAP5 target mRNAs[34].

These cells allowed us to test the effects of eIF4G depletion on the reporter mRNAs in a physiologically relevant context. In the eIF4G depleted cells the cap-dependent translation reporter signal was only 40-50% of the mock treated cells indicating a requirement for eIF4G. The Insr and Igf1r cellular IRES reporters retained more than 80% activity in both a bicistronic and a monocistronic reporter setup, indicating that these IRESes sustain a robust translation activity even when eIF4G1 is depleted. This could indicate that the Insr and Igf1r IRESes utilize DAP5 for activity or that they do not require an eIF4G homologue for initiation.

Taken together these data suggest that the Insr and Igf1r cellular IREes are able to support the initiation of translation independently of the canonical translation factor eIF4G1. They do not rule out, however, that these transcripts may be translated via multiple mechanisms, one of which may be eIF4G1 dependent. When compared to their viral counterparts, the Insr and Igf1r IRESes seem to be more similar to Class III IRESes than Class I or Class II. The HCV IRES typifies this class and requires only eIF3 and a ternary complex in order to initiate translation, but further work on the cellular IRESes will be necessary to determine their precise mechanism.

## Acknowledgements

NKC and MTH were supported for a portion of the work by a training grant from the NIH (T32 GM007122). A portion of the work was supported by a grant to MTM (R01GM117034).

## Author contributions

MTM and NKC conceived the study and wrote the paper. NKC, MTH and MTM performed the experiments. MTM, MTH and NKC analyzed the results and edited and approved the final version of the manuscript.

## Literature Cited

1. Pelletier, J. and N. Sonenberg, The Organizing Principles of Eukaryotic Ribosome Recruitment. Annu Rev Biochem, 2019. 88: p. 307–335.

2. Hiremath, L.S., N.R. Webb, and R.E. Rhoads, Immunological detection of the messenger RNA cap-binding protein. J Biol Chem, 1985. 260(13): p. 7843–9.

3. Duncan, R., S.C. Milburn, and J.W. Hershey, Regulated phosphorylation and low abundance of HeLa cell initiation factor eIF-4F suggest a role in translational control. Heat shock effects on eIF-4F. J Biol Chem, 1987. 262(1): p. 380–8.

4. Ray, B.K., et al., ATP-dependent unwinding of messenger RNA structure by eukaryotic initiation factors. J Biol Chem, 1985. 260(12): p. 7651–8.

5. Pestova, T.V. and V.G. Kolupaeva, The roles of individual eukaryotic translation initiation factors in ribosomal scanning and initiation codon selection. Genes Dev, 2002. 16(22): p. 2906–22.

6. Villa, N., et al., Human eukaryotic initiation factor 4G (eIF4G) protein binds to eIF3c, -d, and -e to promote mRNA recruitment to the ribosome. J Biol Chem, 2013. 288(46): p. 32932–40.

7. LeFebvre, A.K., et al., Translation initiation factor eIF4G-1 binds to eIF3 through the eIF3e subunit. J Biol Chem, 2006. 281(32): p. 22917–32.

8. Korneeva, N.L., et al., Mutually cooperative binding of eukaryotic translation initiation factor (eIF) 3 and eIF4A to human eIF4G-1. J Biol Chem, 2000. 275(52): p. 41369–76.

9. Phan, L., et al., A subcomplex of three eIF3 subunits binds eIF1 and eIF5 and stimulates ribosome binding of mRNA and tRNA(i)Met. EMBO J, 2001. 20(11): p. 2954–65.

10. Siridechadilok, B., et al., Structural roles for human translation factor eIF3 in initiation of protein synthesis. Science, 2005. 310(5753): p. 1513–5.

11. Majumdar, R., A. Bandyopadhyay, and U. Maitra, Mammalian translation initiation factor eIF1 functions with eIF1A and eIF3 in the formation of a stable 40 S preinitiation complex. J Biol Chem, 2003. 278(8): p. 6580–7.

12. De Gregorio, E., T. Preiss, and M.W. Hentze, Translation driven by an eIF4G core domain in vivo. EMBO J, 1999. 18(17): p. 4865–74.

13. Pelletier, J. and N. Sonenberg, Internal initiation of translation of eukaryotic mRNA directed by a sequence derived from poliovirus RNA. Nature, 1988. 334(6180): p. 320–5.

14. Sonenberg, N. and J. Pelletier, Poliovirus translation: a paradigm for a novel initiation mechanism. Bioessays, 1989. 11(5): p. 128–32.

15. Holcik, M. and N. Sonenberg, Translational control in stress and apoptosis. Nat Rev Mol Cell Biol, 2005. 6(4): p. 318–27.

16. Balvay, L., et al., Structural and functional diversity of viral IRESes. Biochim Biophys Acta, 2009. 1789(9-10): p. 542–57.

17. Fernandez-Miragall, O., S. Lopez de Quinto, and E. Martinez-Salas, Relevance of RNA structure for the activity of picornavirus IRES elements. Virus Res, 2009. 139(2): p. 172–82.

18. Kwan, T. and S.R. Thompson, Noncanonical Translation Initiation in Eukaryotes. Cold Spring Harb Perspect Biol, 2019. 11(4).

19. Jackson, R.J., The current status of vertebrate cellular mRNA IRESs. Cold Spring Harb Perspect Biol, 2013. 5(2).

20. Olson, C.M., et al., The insulin receptor cellular IRES confers resistance to eIF4A inhibition. eLife, 2013. 2(300542).

21. Marr II, M.T., et al., IRES-mediated functional coupling of transcription and translation amplifies insulin receptor feedback. Genes & Development, 2007. 21: p. 175–183.

22. Vasudevan, D., et al., The GCN2-ATF4 Signaling Pathway Induces 4E-BP to Bias Translation and Boost Antimicrobial Peptide Synthesis in Response to Bacterial Infection. Cell Rep, 2017. 21(8): p. 2039–2047.

23. Lindqvist, L., et al., Selective pharmacological targeting of a DEAD box RNA helicase. PLoS One, 2008. 3(2): p. e1583.

24. Nabet, B., et al., The dTAG system for immediate and target-specific protein degradation. Nat Chem Biol, 2018. 14(5): p. 431–441.

25. David, A., et al., Nuclear translation visualized by ribosome-bound nascent chain puromycylation. J Cell Biol, 2012. 197(1): p. 45–57.

26. Ventoso, I., et al., Poliovirus 2A proteinase cleaves directly the eIF-4G subunit of eIF-4F complex. FEBS Lett, 1998. 435(1): p. 79–83.

27. Svitkin, Y.V., et al., Eukaryotic translation initiation factor 4E availability controls the switch between cap-dependent and internal ribosomal entry site-mediated translation. Mol Cell Biol, 2005. 25(23): p. 10556–65.

28. Hinton, T.M., et al., Functional analysis of individual binding activities of the scaffold protein eIF4G. J Biol Chem, 2007. 282(3): p. 1695–708.

29. Nabet, B., et al., Rapid and direct control of target protein levels with VHL-recruiting dTAG molecules. Nat Commun, 2020. 11(1): p. 4687.

30. Nabet, B., et al., Rapid and direct control of target protein levels with VHL-recruiting dTAG molecules. Nature Communications, 2020. 11(4687).

31. Ingolia, N.T., L.F. Lareau, and J.S. Weissman, Ribosome profiling of mouse embryonic stem cells reveals the complexity and dynamics of mammalian proteomes. Cell, 2011. 147(4): p. 789–802.

32. Witherell, G.W. and E. Wimmer, Encephalomyocarditis virus internal ribosomal entry site RNA-protein interactions. J Virol, 1994. 68(5): p. 3183–92.

33. Ramirez-Valle, F., et al., eIF4GI links nutrient sensing by mTOR to cell proliferation and inhibition of autophagy. J Cell Biol, 2008. 181(2): p. 293–307.

34. de la Parra, C., et al., A widespread alternate form of cap-dependent mRNA translation initiation. Nat Commun, 2018. 9(1): p. 3068.

